## Supplementary Materials for "Identification of Druggable Binding Sites and Small Molecules as Modulators of TMC1"

[*]Correspondence should be addressed to:

[&]Current address:

**Supplementary Table 1.** Known MET channel blockers used to build a common pharmacophore model (related to *Figure 1*)

| ID | Compound | Concentration reported | MET channel | Reference |
|---|---|---|---|---|
| **1** | UoS-7692 | 50 µM | 80% block | (Kenyon et al., 2021) |
| **2** | UoS-3607 | 50 µM | 70% block | (Kenyon et al., 2021) |
| **3** | UoS-3606 | 50 µM | 80% block | (Kenyon et al., 2021) |
| **4** | UoS-5247* | 50 µM | 20% block (Steep release) | (Kenyon et al., 2021) |
| **5** | UoS-962 | 50 µM | 20% block | (Kenyon et al., 2021) |
| **6** | Proto-1 | 10 µM | 80% control hair cells | (Owens et al., 2008) |
| **7** | E6 berbamine | 25 µM | Significant hair cells damage | (Kruger et al., 2016) |
| **8** | Hexamethyleneamiloride | 10 µM | 50% control hair cells survival | (Ou et al., 2009) |
| **9** | Amsacrine | 10 µM | 50% control hair cells survival | (Ou et al., 2009) |
| **10** | Phenoxybenzamine | 10 µM | 39% control hair cells survival | (Ou et al., 2009; Majumder et al., 2017) |
| **11** | Carvedilol derivate 13 | 5 µM | 50% block | (O'Reilly et al., 2019) |
| **12** | ORC-13661 | 8.3 µM | 85% control hair cells | (Kitcher et al., 2019; Sarwat Chowdhury, et al., 2018) |
| **13** | FM1-43 | 20 µM | 95% block | (Gale et al., 2001, Jason R Meyers et al., 2003) |

* UoS-5247 has been reported as a week MET blocker (Kenyon et al., 2021). We included this compound to introduce structural diversity due its lipophilic substituent (See also *Figure 1*)

**Supplementary Table 2.** Matching compounds for each pharmacophore. Compounds that match the pharmacophore are marked with an X. Hydrogen bond acceptor (*A*), positively charged group (*P*), aromatic ring (*R*), and hydrophobic group (*H*).

| Compound | APRR | APR-1 | APR-2 | PRR | ARR-1 | ARR-2 | AHR | ARR-3 | ARR-4 | HRR | Total matches |
|---|---|---|---|---|---|---|---|---|---|---|---|
| **UoS-7692** | X | X | X | X | X | X | X | | X | X | 9 |
| **UoS-3607** | X | X | X | X | X | X | X | X | X | | 9 |
| **UoS-3606** | | | | | X | | X | | | | 2 |
| **UoS-5247 *** | | | | | | | | | | | 0 |
| **UoS-962** | X | X | X | X | X | X | | | X | X | 8 |
| **Proto-1** | | | | | | X | | X | X | X | 4 |
| **E6 bermamine** | **X** | **X** | **X** | **X** | **X** | **X** | **X** | **X** | **X** | **X** | **10** |
| **Hexamethyleneamiloride** | | X | | | | | X | | | | 2 |
| **Amsacrine** | X | X | X | X | | X | | X | | X | 7 |
| **Phenoxybenzamine** | **X** | **X** | **X** | **X** | **X** | **X** | **X** | **X** | **X** | **X** | **10** |
| **Carvedilol derivate 13** | **X** | **X** | **X** | **X** | **X** | **X** | **X** | **X** | **X** | **X** | **10** |
| **ORC-13661** | | X | X | | | X | | X | X | X | 6 |
| **FM1-43** | | | | | | | | | | X | 1 |
| **Total** | 7 | 9 | 8 | 7 | 7 | 9 | 7 | 7 | 8 | 9 | |

***** UoS-5247 is a weak MET blocker previously reported to have a steep release activity (Kenyon et al., 2021). It was selected to increase structural diversity due its lipophilic substituent. However, upon Phase analysis, UoS-5247 did not fit to any of the ten pharmacophores used in this study, suggesting that the lipophilic substituent and the absence of a bigger aromatic moiety, such as benzimidazole (also present in UoS-7692, UoS-3607, UoS-3606), could be detrimental to the interaction with the MET channel (Kenyon et al., 2021).

**Supplementary Table 3.** Pharmacophore model validation using Güner-Henry (GH) scoring method.

| | APRR | APR-1 | APR-2 | PRR | ARR-1 | ARR-2 | AHR | ARR-3 | ARR-4 | HRR |
|---|---|---|---|---|---|---|---|---|---|---|
| **Total actives** | 7 | 9 | 8 | 7 | 7 | 9 | 7 | 7 | 8 | 9 |
| **Decoys** | 280 | 360 | 320 | 280 | 280 | 360 | 280 | 280 | 320 | 360 |
| **Actives+Decoys** | 287 | 369 | 328 | 287 | 287 | 369 | 287 | 287 | 328 | 369 |
| **True positives (TP)** | 7 | 8 | 8 | 7 | 7 | 9 | 7 | 7 | 8 | 9 |
| **False positive (FP)** | 2 | 4 | 4 | 4 | 5 | 5 | 4 | 4 | 5 | 6 |
| **True negative (TN)** | 278 | 356 | 316 | 276 | 275 | 355 | 276 | 276 | 315 | 354 |
| **False negative (FN)** | 0 | 1 | 0 | 0 | 0 | 0 | 0 | 0 | 0 | 1 |
| **ROC (AUC)** | 1 | 0.86 | 0.99 | 0.99 | 0.98 | 0.99 | 0.98 | 0.99 | 0.99 | 0.99 |
| **Sensitivity ($S_e$)** | 1 | 0.889 | 1 | 1 | 1 | 1 | 1 | 1 | 1 | 0.900 |
| **Specificity ($S_p$)** | 0.993 | 0.989 | 0.988 | 0.986 | 0.982 | 0.986 | 0.986 | 0.986 | 0.984 | 0.983 |
| **% Yield of active compounds ($Y_a$)** | 78 | 67 | 67 | 64 | 58 | 64 | 64 | 64 | 62 | 60 |
| **Enrichment factor (EF)** | 31.88 | 27.333 | 27.333 | 26.09 | 23.917 | 26.357 | 26.091 | 26.091 | 25.231 | 24.600 |
| **% Yield** | 100 | 88.889 | 100 | 100 | 100 | 100 | 100 | 100 | 100 | 100 |
| **Güner–Henry (GH) scoring** | 0.827 | 0.714 | 0.741 | 0.717 | 0.675 | 0.722 | 0.717 | 0.717 | 0.700 | 0.664 |

**A:** Hydrogen-bond acceptor. **P:** positively charged group. **R:** aromatic ring. **H:** hydrophobic group. **ROC:** Receiver Operating Characteristic. **AUC:** Area Under the ROC Curve.

**Supplementary Table 4.** Matrix constructed with the *PhaseScreenScore* that each compound of the non-FDA-approved *Library* 1 obtained for each pharmacophore after 2nd-step PBVS with *Phase*. The top-5 hits (according to the *TPSₛ*, as well as the last 5 solutions are presented

| N° | Compound | APRR | APR-1 | APR-2 | PRR | ARR-1 | ARR-2 | AHR | ARR-3 | ARR-4 | HRR | TPSs |
|---|---|---|---|---|---|---|---|---|---|---|---|---|
| 1 | ZINC26876007 | 2.449 | 2.272 | 2.290 | 2.438 | 2.103 | 2.208 | 2.080 | 1.708 | 2.054 | 1.910 | 21.512 |
| 2 | ZINC93401109 | 2.045 | 2.298 | 2.357 | 2.309 | 1.994 | 2.203 | 1.700 | 1.778 | 2.165 | 2.152 | 21.001 |
| 3 | ZINC51125638 | 2.172 | 2.307 | 2.081 | 2.323 | 1.998 | 2.209 | 1.958 | 2.043 | 1.945 | 1.884 | 20.920 |
| 4 | ZINC91304884 | 2.439 | 2.407 | 2.287 | 2.378 | 2.150 | 2.317 | 1.668 | 1.676 | 1.895 | 1.700 | 20.917 |
| 5 | ZINC60557463 | 2.094 | 2.274 | 2.241 | 2.243 | 2.015 | 2.341 | 1.653 | 1.755 | 2.111 | 2.065 | 20.792 |
| … | … | … | … | … | … | … | … | … | … | … | … | … |
| 4183 | ZINC39970059 | 0 | 0 | 0 | 0 | 1,506 | 0 | 0 | 0 | 0 | 0 | 1,506 |
| 4184 | ZINC39967993 | 0 | 0 | 0 | 0 | 1,337 | 0 | 0 | 0 | 0 | 0 | 1,337 |
| 4185 | ZINC63373196 | 0 | 0 | 0 | 0 | 1,309 | 0 | 0 | 0 | 0 | 0 | 1,309 |
| 4186 | ZINC39708296 | 0 | 0 | 0 | 0 | 1,245 | 0 | 0 | 0 | 0 | 0 | 1,245 |
| 4187 | ZINC39969398 | 0 | 0 | 0 | 0 | 0 | 0 | 1.157 | 0 | 0 | 0 | 1.157 |

**A**: hydrogen-bond acceptor. **P**: positively charged group. **R**: aromatic ring. **H**: hydrophobic group. ***TPSs***: *Total-PhaseScreenScore*

**Supplementary Table 5.** Matrix built with the *PhaseScreenScore* that each compound of the FDA-approved drug (*Library* 2) obtained for each pharmacophore after 2nd-step PBVS with *Phase*. The top-5 hits (according to the *TPSs*, as well as the last 5 solutions are presented

| N° | Compound | APRR | APR-1 | APR-2 | PRR | ARR-1 | ARR-2 | AHR | ARR-3 | ARR-4 | HRR | TPSs |
|---|---|---|---|---|---|---|---|---|---|---|---|---|
| 1 | Carvedilol | 1.736 | 2.301 | 1.762 | 1.666 | 2.298 | 2.168 | 1.651 | 1.544 | 1.825 | 1.642 | 18.593 |
| 2 | Candesartan Cilexetil | 1.498 | 1.609 | 1.465 | 1.776 | 1.981 | 1.923 | 1.912 | 1.809 | 2.079 | 1.698 | 17.750 |
| 3 | Olmesartan Medoxomil | 1.532 | 1.511 | 1.508 | 1.557 | 2.212 | 1.999 | 1.719 | 1.886 | 1.968 | 1.767 | 17.659 |
| 4 | Candesartan | 1.602 | 1.709 | 1.532 | 1.759 | 1.821 | 2.041 | 1.964 | 1.599 | 1.895 | 1.551 | 17.473 |
| 5 | Amitraz | 1.836 | 2.200 | 1.297 | 1.924 | 1.513 | 1.867 | 1.985 | 1.486 | 1.822 | 1.484 | 17.414 |
| … | … | … | … | … | … | … | … | … | … | … | … | … |
| 902 | Nortrupityline | 0 | 0 | 0 | 0 | 0 | 0,636 | 0 | 0 | 0 | 0 | 0,636 |
| 903 | Estradiol Cypionate | 0,62 | 0 | 0 | 0 | 0 | 0 | 0 | 0 | 0 | 0 | 0,620 |
| 904 | Valganciclovir | 0 | 0,547 | 0 | 0 | 0 | 0 | 0 | 0 | 0 | 0 | 0,547 |
| 905 | Cefcapene Pivoxil | 0,506 | 0 | 0 | 0 | 0 | 0 | 0 | 0 | 0 | 0 | 0,506 |
| 906 | Trimeprazine | 0 | 0 | 0 | 0 | 0 | 0 | 0.419 | 0 | 0 | 0 | 0.419 |

**A:** hydrogen-bond acceptor**. P:** positively charged group**. R:** aromatic ring**. H:** hydrophobic group**.** ***TPSs***: *Total-PhaseScreenScore*

**Supplementary Table 6.** List of key residues located in the top, middle, and bottom binding sites of TMC1 involved in binding of small molecules (related to colored regions in *Figure 4* and *Figure 5*)

| Sites | Residues |
|---|---|
| Top | N404, M407, F451, E458, E520, R523, L524, S527, M583, POPC-B |
| Middle | S408, **G411**, **M412**, L443, L444, **N447**, L448, **D528**, T531, **T532**, I536, F579, N580, G582, R601 |
| Bottom | Y182, P415, T416, D419, L436, L437, I440, T535, D540, R543, T566, E567, F568, **D569**, S571, **G572**, N573, L575, A576, L577, Q609, A612, V613, N617, POPC-A |
| Zwitterionic-like network ZN1* | D239, N404, E458, E520, R523, POPC-B |
| Zwitterionic-like network ZN2 | **D528**, T531, **T532**, N580, N598, R601 |
| Zwitterionic-like network ZN3* | T416, D419, E423, T535, D540, R543, D569, N573, E620, R622 |

*The polar head of POPC-A points towards the bottom site forming a zwitterionic-like interaction network with R543, D569, E567, E620, and R622, key for both ligand and phospholipid binding. Additionally, the polar head of POPC-B forms a similar zwitterionic-like network with R523, N401, and N404 residues. Hydrophobic tails of both phospholipids form a sidewall between TM4 and TM6 (see **Supp Figure 4**). Residues in bold are known deafness mutation sites or known to alter TMC1 leak courrents[1].

| Supplementary Table 7. Re-scoring of docking poses for 16 known blockers using MM-GBSA. | | |
|---|---|---|
| Compound | Without phospholipids $\Delta G_{bind}$ (kcal/mol) | With POPC-B and POPC-A $\Delta G_{bind}$ (kcal/mol) |
| E6-berbamine | -73.65 | -82.72 |
| FM1-43 | -57.69 | -84.44 |
| UoS-3607 | -54.80 | -61.99 |
| Carvedilol derivate 13 | -45.35 | -53.44 |
| Tubocurarine(*R*) | -44.32 | -44.79 |
| Hexamethyleneamiloride | -43.56 | -42.01 |
| UoS-962 | -41.48 | -71.15 |
| UoS-5247 | -34.99 | -51.17 |
| UoS-3606 | -33.38 | -40.21 |
| Phenoxybenzamine(*R*) | -33.26 | -36.99 |
| UoS-7692 | -31.34 | -23.79 |
| Proto-1 | -29.81 | -37.71 |
| ORC-13661 | -27.76 | -40.87 |
| Amsacrine | -24.16 | -49.74 |
| Benzamil | -20.96 | -36.48 |
| Phenoxybenzamine(*S*) | -19.38 | -43.06 |

**Supplementary Table 8.** Structural diversity analysis results for both *Library* 1 and *Library* 2 ($\Delta G_{bind}$ = kcal/mol) without phospholipids.

| Library 1 (non-FDA-approved compounds) | | | Library 2 (FDA-approved drugs) | | |
|---|---|---|---|---|---|
| Cluster ID | Population (# compounds) | Average $\Delta G_{bind}$ ± SD | Cluster ID | Population (# compounds) | Average $\Delta G_{bind}$ ± SD |
| 1.1 | 11 (3)* | -24.74 ± 5.50 | 2.1 | 17 (1)* | -34.49 ± 6.46 |
| 1.2 | 7 (3)* | -33.58 ± 10.81 | 2.2 | 10 (1)* | -34.46 ± 9.82 |
| 1.3 | 4 (1)* | -31.65 ± 4.81 | 2.3 | 6 (1)* | -36.38 ± 6.57 |
| 1.4 | 4 (1)* | -37.62 ± 37.62 | 2.4 | 5 (1)* | -21.98 ± 7.57 |
| 1.5 | 3 (1)* | -20.18 ± 2.01 | 2.5 | 4 (1)* | -45.88 ± 16.03 |
| 1.6 | 3 (1)* | -39.04 ± 11.70 | 2.6 | 3 (1)* | -38.85 ± 14.44 |
| 1.7 | 2 | -25.18 ± 4.55 | 2.7 | 2 (1)* | -33.68 ± 8.31 |
| 1.8 | 2 | -36.65 ± 19.03 | 2.8 | 2 (1)* | -25.05 ± 0.54 |
| 1.9 | 2 | -21.55 ± 4.99 | 2.9 | 1 (1)* | -24.10 |
| 1.10 | 2 | -30.21 ± 1.02 | 2.10 | 1 (1)* | -20.68 |
| 1.11 | 1 | -26.81 | | | |
| 1.12 | 1 | -37.15 | | | |
| 1.13 | 1 | -24.29 | | | |
| 1.14 | 1 | -26.47 | | | |
| 1.15 | 1 | -27.16 | | | |
| Total = 15 | Total = 45 (10)* | | Total = 10 | Total = 51 (10)* | |

*Number of compounds with best $\Delta G_{bind}$ suitable for experimental evaluation

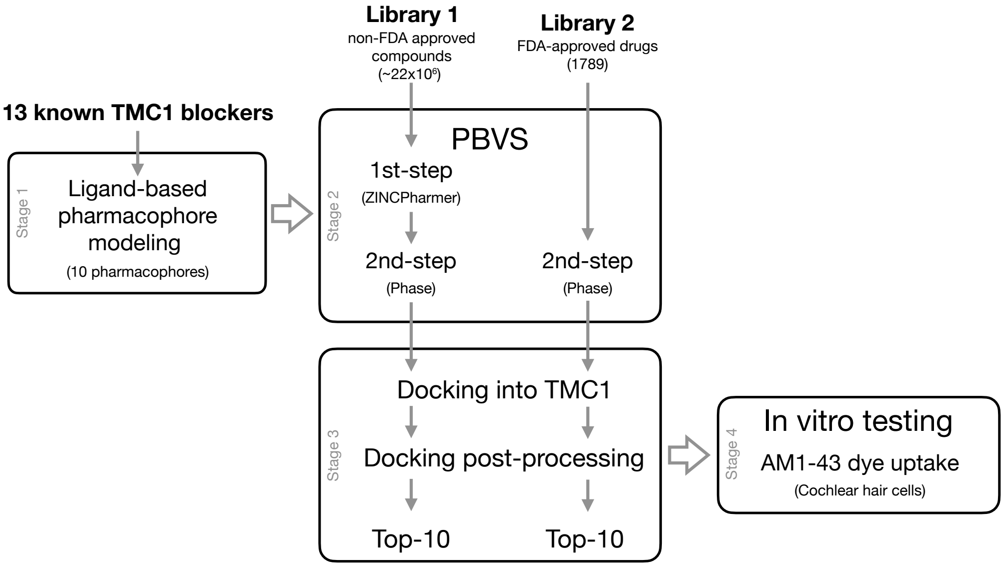


**Supplementary Figure 1.** Systematic workflow employed to identify TMC1 modulators. Initially, 10 commercially available top hit compounds were selected from each of two libraries: *Library* 1 (ZINC Database)[93] and *Library* 2 (MicroSource Discovery Systems)[94] resulting in a total of 20 hit compounds (see *Methods* for details).

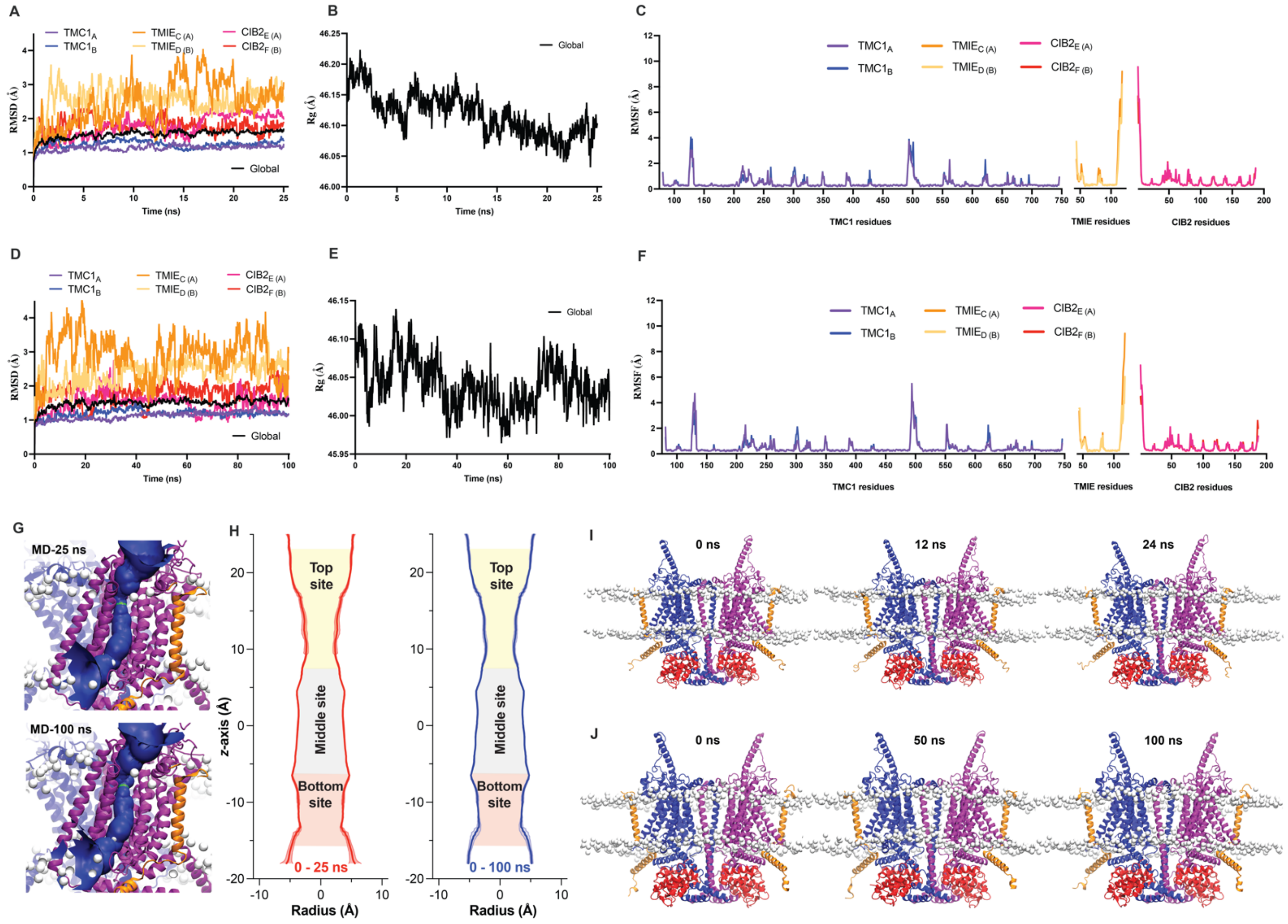


**Supplementary Figure 2. Molecular dynamics simulations of MET complex. (A)** Time dependence of the Root Mean Squared Deviation (RMSD) for TMC1, TMIE and CIB2 backbones during the 25ns MDs. **(B)** Radius of gyration (*Rg*) to study the compactness of the protein. **(C)** Root Mean Squared Fluctuation (RMSF) characterizing the internal residue fluctuation of TMC1, TMIE, and CIB2 during the 25 ns MDs. **(D)** Time dependence of the RMSD for TMC1, TMIE and CIB2 backbones during the 100 ns MDs. **(E)** *Rg* to study the compactness of the protein as in panel *b*. **(F)** RMSF characterizing the internal residue fluctuation of TMC1, TMIE and CIB2 during the 100 ns MDs. **(G)** TMC1 pore of chain A represented by a blue funnel obtained by HOLE after 25 ns and 100 ns. **(H)** Pore radius measurements (average ± standard deviation) of the TMC1 chain A obtained by HOLE analysis over 25 ns and 100 ns simulation trajectories. Measurements were obtained every nanosecond throughout the simulations. The top, middle, and bottom sites of the pore are color-coded as in *Figure 2B* and *Figure 4*. The zero point (0 Å) on the z-axis represents the reference position of the middle site of the TMC1 pore at the center of the plasma membrane. **(I-J)** General view and structural comparison of three representative frames from the 25 ns and 100 ns MD simulations, respectively, revealing no major structural changes between the simulations time points and suggesting stability across the trajectories.

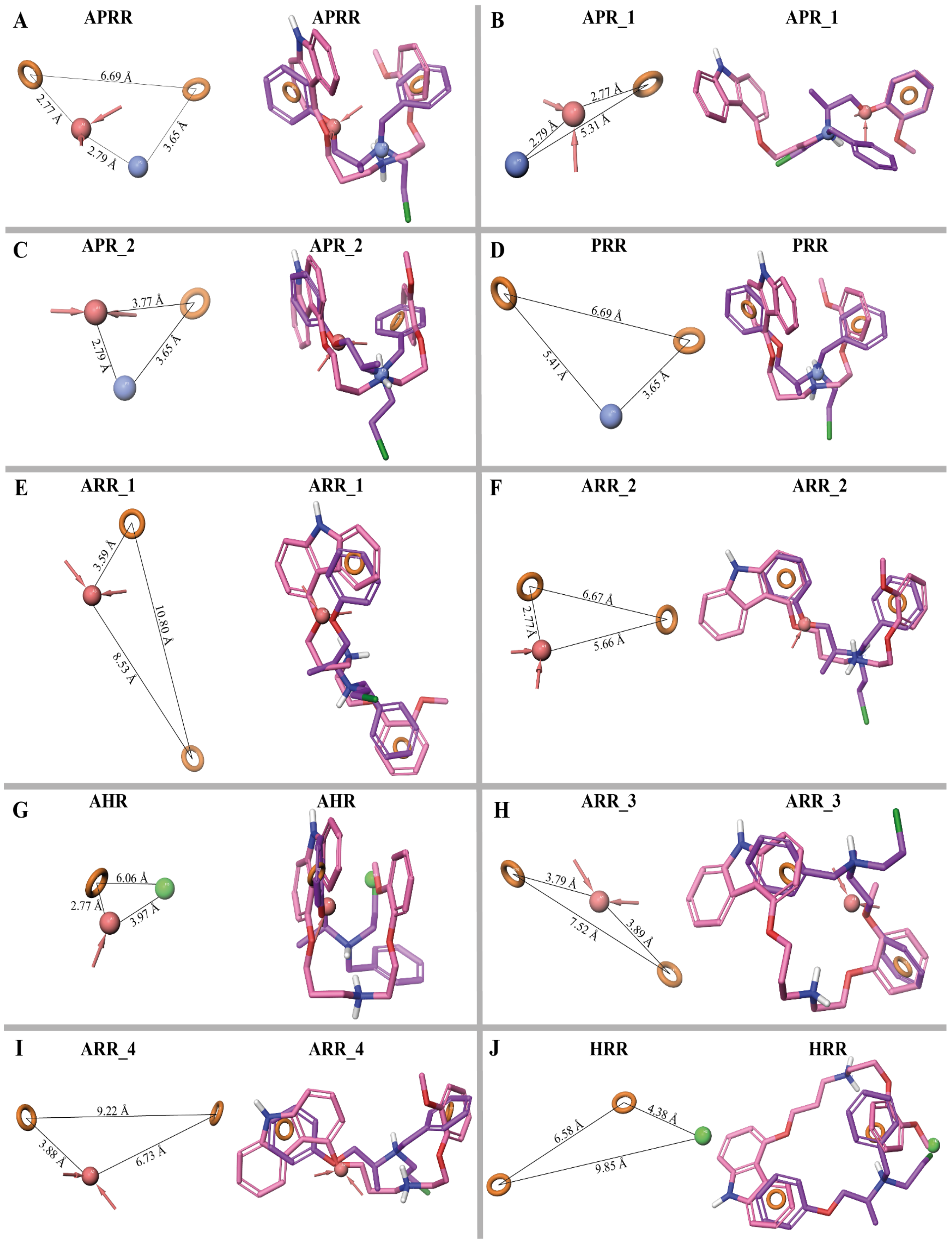


**Supplementary Figure 3. 3D view of the 10 common pharmacophores of 13 known MET channel blockers**. **(A-K)** Pharmacophore models of MET channel blockers (see *Figure 1*). Hydrogen-bond acceptor features (*A*) in red, aromatic rings (*R*) in orange circles, positively charged group (*P*) in blue, and hydrophobic group (*H*) in green beads. Distances between the vector features are labeled. Superposed structures of the MET blockers phenoxybenzamine (purple) and carvedilol derivative 13 (pink) with the vector features of each pharmacophore. These two compounds fit to all the 10 pharmacophores reported in Table 1 (See *Supp Table 2*). 3D structures were depicted with Pymol[2].

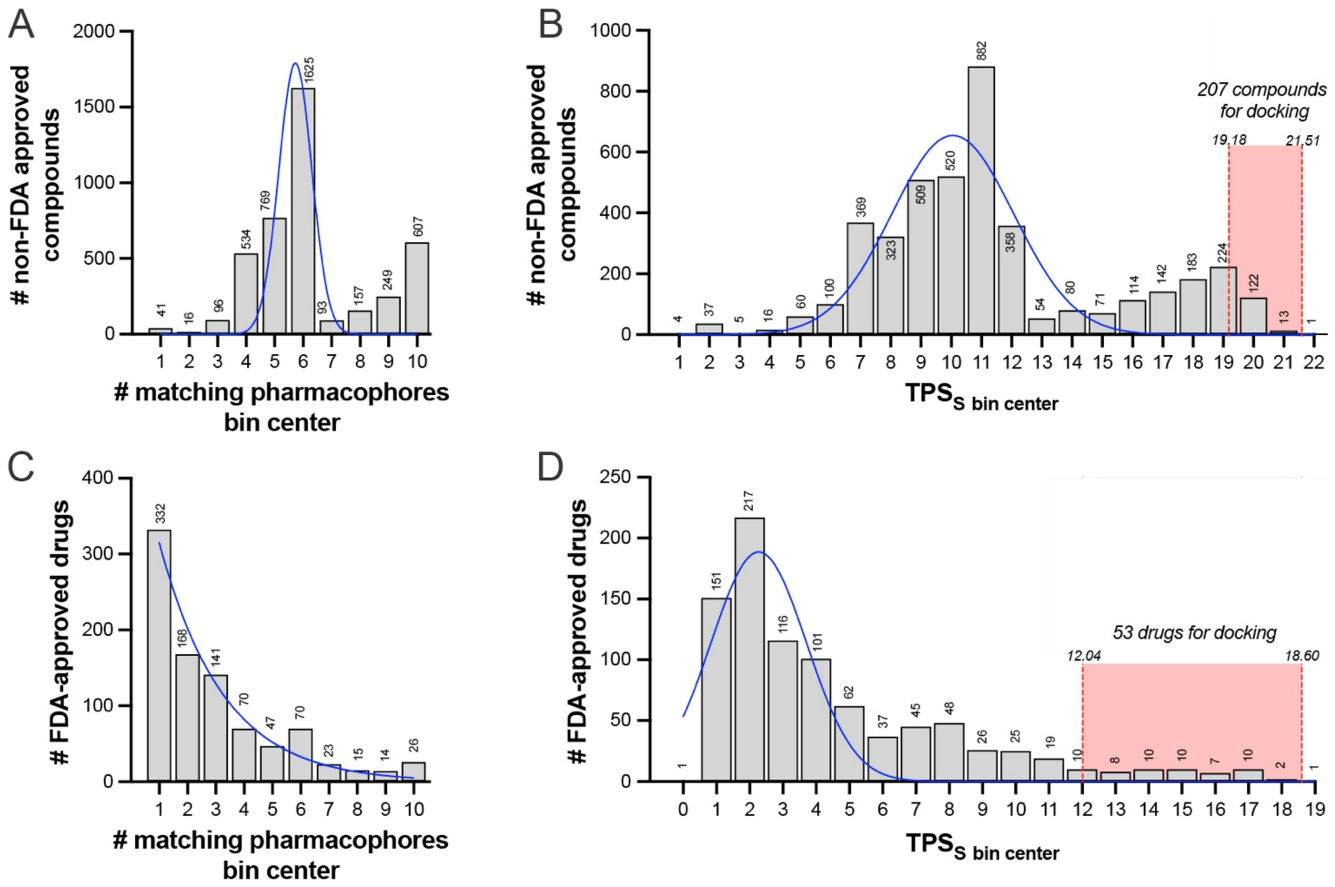


**Supplementary Figure 4. Second step PBVS analysis results for Libraries 1 and 2. (A)** Histogram reporting the number of compounds presented as a function of the number of pharmacophores they matched following the second step PBVS in *Phase*. **(B)** Frequency distribution of non-FDA-approved compounds from *Library* 1 as a function of their *Total-PhaseScreenScore* (*$TPS_S$*). Red shaded area represents the compounds with *$TPS_S$* score values above the cut-off docking threshold (Mean + 2 SD) as a selection criterion. **(C)** Histogram reporting the number of FDA-approved compounds presented as a function of the number of pharmacophores they matched following PVBS in *Phase*. **(D)** Frequency distribution of FDA-approved drugs from *Library* 2 as a function of their *TPSs*. Red shaded area represents the compounds with *$TPS_S$* score values above the cut-off docking threshold (Mean + 2 SD) as a selection criterion. Values above the bars report the number of compounds per bin. Blue lines represent the data fit for normal (Gaussian) distribution.

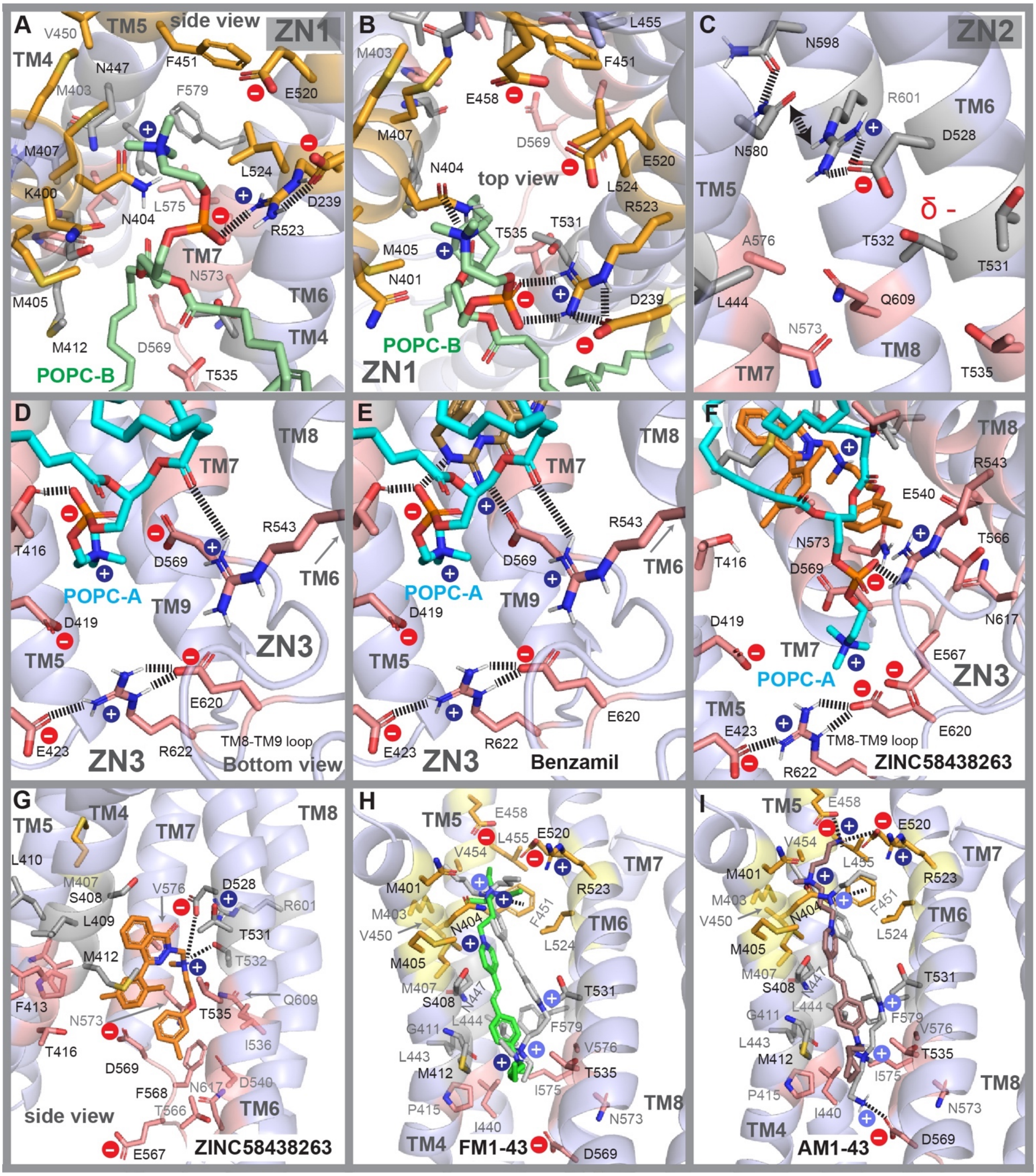


**Supplementary Figure 5. Zwitterionic sites and binding modes of ZINC58438263, FM1-43 and AM143 in the TMC1 pore. (A-B)** Side and top views of the ZN1 site showing POPC-B (lime). Dashed lines represent interactions between the negative head of the phospholipid with R523 and D239. The positive charged iminium group is placed near N404 forming salt bridges. **(C)** Side view of the ZN2 site. TM4 was removed for clarity. The amino acids N580, N598, R601, and D528 closely interact forming hydrogen bond and salt bridge interactions, stabilizing TM5, TM6, and TM8 at the middle site of the pore. The residues D528, T531, and T532 form an electronegatively charged area ($\delta^-$) involving in ligand binding. **(D-E)** Bottom view of the ZN3 site in absence and presence of benzamil. Benzamil displays key interactions with D569 and the negative head of POPC-A, which in turn interact with T416 forming hydrogen bond interactions with TM4. Similarly, the carbonyl group of POPC-A interacts with R543 located in TM6. The ZN3 site might be relevant for phospholipid and ligand stabilization at the bottom site. Furthermore, other stabilizing interactions are displayed between R622 and the charged residues E423 and E620 which clamp TM4, TM8, and TM9 with salt-bridge interactions, stabilizing the bottom cavity. **(F)** Bottom view of the ZN3 site and binding of ZINC58438263 along with POPC-A. ZINC58438263 does not displays direct interactions with D569 as benzamil. Interactions between R622, E423, and E620 are also showed as in panel C-E. **(G)** Side view of the binding mode of ZINC58438263, displaying direct interactions between the iminium group and both D528 and T532. Neighboring residues are showed, specially M412 forming van der Waals interactions. **(H)** Two binding docking conformations predicted for FM1-43. The first binding mode conformation of FM1-43 (green) shows its triethylammonium charged group forming a cation-$\pi$ stacking interaction with F451 at the top site near ZN1. The second binding mode (in gray) is inverted pointing the triethylammonium charged group towards the bottom site near M412 and T535. In this scenario, the positively charged dibutylamine group of FM1-43 is not placed near F451 as analyzed above. **(I)** Side view of the binding mode of AM1-43 (brown). This first binding mode conformation shows similar interactions to those displayed by FM1-43 (panel H, green), with the difference that the elongated and protonated butylamine group forms interactions with E458 and E520 at the ZN1 site near the entry of the pore. The second binding mode of AM1-43 (gray) is inverted with the same group now forming a direct salt bridge with D569 at the bottom site. In this second scenario, the positively charged dibutylamine group, mimics the same cation-$\pi$ stacking interaction with F451 displayed by the triethylammonium charged group of FM1-43, as in panel H.

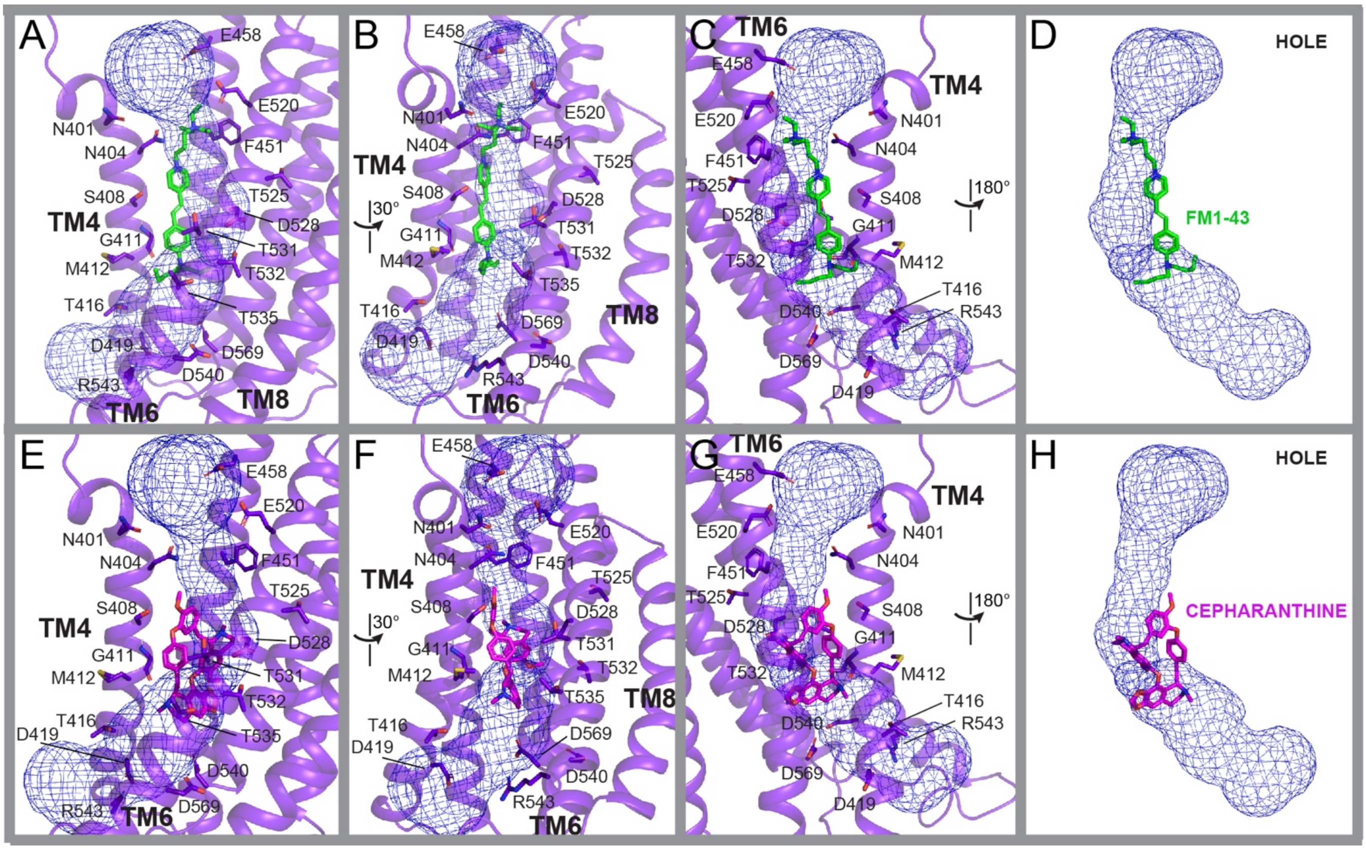


**Supplementary Figure 6. Structural features of TMC1 pore with HOLE including ligands. (A-D)** Structural representation of the pore and FM1-43. **(A)** Side view of a TMC1 protomer showing a curved pore calculated by HOLE[3] in blue mesh representation. Docking pose of FM1-43 is showed in green licorice representation (as in *Figure 5* and *6*). **(B-C)** 30° and 180° rotation view from panel *a*, respectively. **(B)** View of the mesh contour and FM1-43 as in panel *c***.** As predicted by docking, some areas of the FM1-43 molecule are outside of the mesh contour predicted by HOLE, suggesting that some ligand binding-residues are different than those used by HOLE to calculate the mesh-pore contour. **(E-H)** Structural representation of the pore and the docked cepharanthine, as showed in panels *a-d*. Similar to FM1-43, a portion of the cepharanthine molecule is docked inside the mesh contour predicted by HOLE, while two aromatic moieties are placed outside the mesh contour. In both cases, the positively charged nitrogen atoms of FM1-43 and cepharanthine are located inside the mesh contour of the pore. Results suggest that both elongated and spherical ligands bind to the TMC1 pore occupying regions not predicted by HOLE.

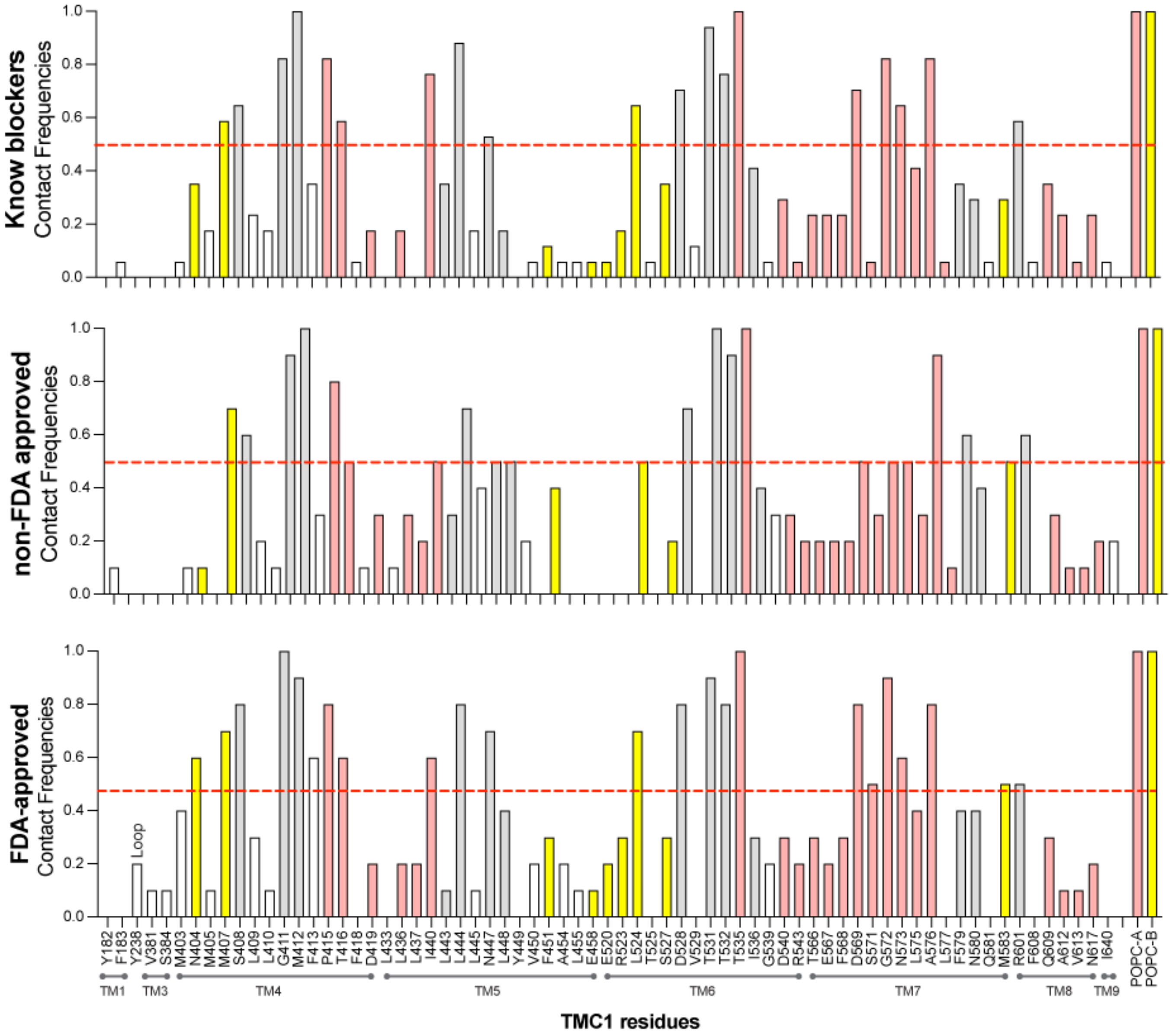


**Supplementary Figure 7. Contact frequencies of residues involved in TMC1-ligand binding across 36 analyzed compounds.** Contact frequencies, representing the portion of ligands interacting with the respective residue. Frequency columns are colored according to the location of the residue within the top (*gold*), middle (*light gray*), and bottom (*light red*) sites of the TMC1 pore. White columns indicate residues not identified within the mentioned pore sites. Red dashed line indicates a contact frequency of 0.5 (50%). *Top panel,* contact frequencies between TMC1 residues and the known blockers; *Middle panel,* contact frequencies between TMC1 residues and the hit compounds from *Library* 1; *Bottom panel,* contact frequencies between TMC1 residues and the hit drugs from *Library* 2. TM domains and the respective residues are labeled at the bottom of the figure.

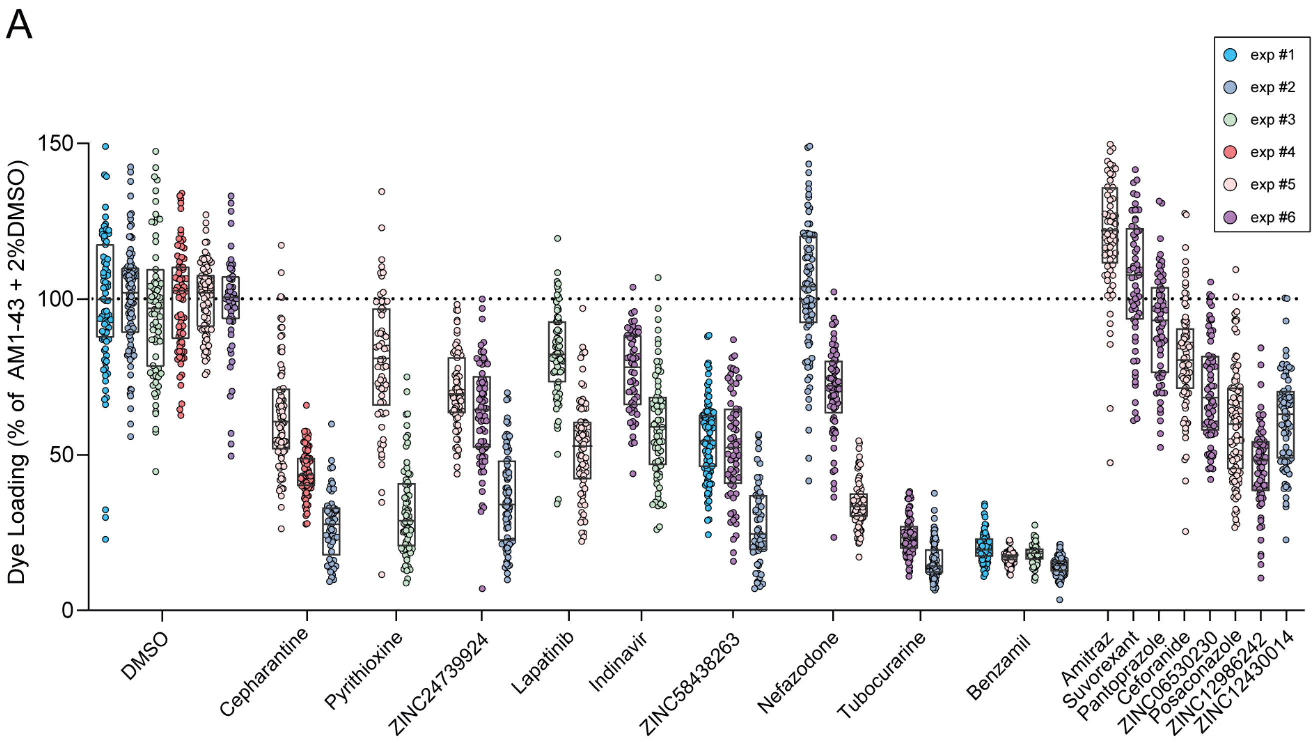


**Supplementary Figure 8. Quantification of AM1-43 dye uptake by individual OHCs across all experimental sessions for both compounds and controls.** Each dot represents the fluorescence intensity of an individual OHC normalized to the average fluorescence of the DMSO controls from the same experimental session. The boxes represent the interquartile range, spanning from the 25th percentile to the 75th percentile. The line inside the box indicates the median.

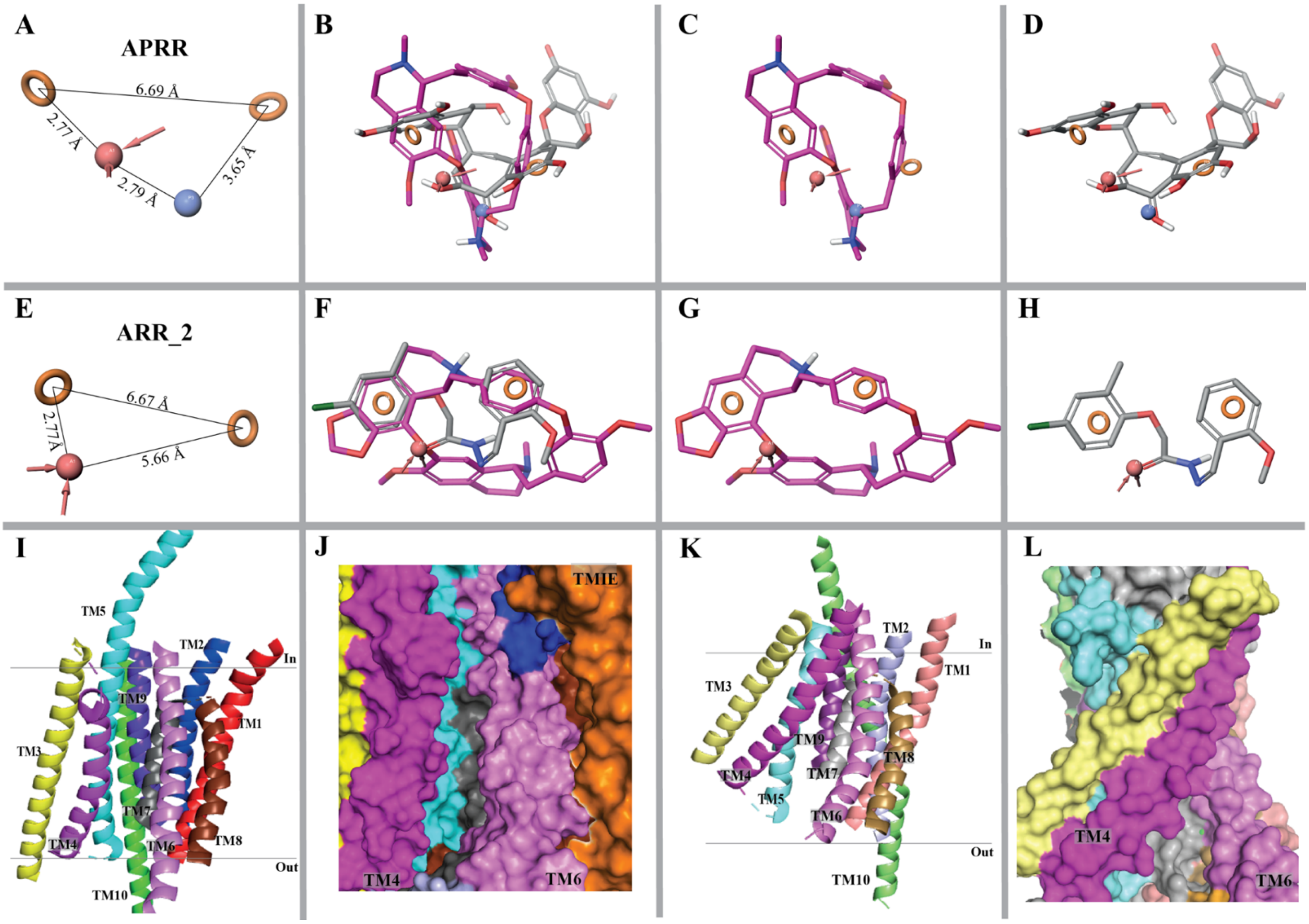


**Supplementary Figure 9. 3D view of the 2 common pharmacophores of TMC1 and TMEM16 channel blockers**. **(A** and **E)** Pharmacophore models of MET channel blockers (see *Figure 1*), ID1 and ID6, respectively (see *Table1*). **(A)** APRR pharmacophore model. Hydrogen-bond acceptor features (*A*) in red, aromatic rings (*R*) in orange circles and positively charged group (*P*) in blue beads. Distances between the vector features are labeled. **(B)** Superposed structures of the hit compound cepharanthine (magenta) and the theaflavin with a protonated carbonyl group (gray) with the vector features of the TMC1-pharmacophore APRR. **(C-D)** Structural alignment of cepharanthine and theaflavin with the APRR vector features, respectively. **(E)** ARR_2 pharmacophore model. **(F)** Superposed structures of the hit compound cepharanthine (magenta) and Ani9 (gray) with the vector features of the TMC1-pharmacophore ARR-2. **(G-H)** Structural alignment of cepharanthine and Ani9 with the ARR-2 vector features, respectively. Theaflavin and Ani9 are inhibitors of TMEM16A[4]. **(I)** Cartoon representation of an open-like conformation of TMC1. Loops are removed for clarify and TM domains are colored and labeled for clarity. TM4 is colored in magenta and TM6 is colored in dark pink as a reference, showing a parallel (I I) conformation. **(J)** Surface representation of TMC1 protomer as in *Figure 5*. TM4 and TM6 are colored as in panel I*,* showing the parallel conformation. Other TM domains are colored as in panel I, along with TMIE as a reference (dark orange). **(K)** Cartoon representation of TMEM16A as in panel I. TM4 is colored in light magenta and TM6 in light pink, both showing an inverted V conformation clashing at the upper-top site. **(L)** Surface representation of TMEM16A. View and colors as in panel J. TM4 (light magenta) and TM6 (light pink) forms a V conformation that closes the pore at the upper-top site. A space cavity (cyan, gray, and light brown) is shown at the bottom between the separated TM4 and TM6 domains.
